# The absence of collagen VI reduces systolic function but paradoxically increases Ca^2+^ release in the rat heart

**DOI:** 10.1101/2025.03.21.644665

**Authors:** A Krstic, S Hassan, H Moammer, J Bai, P Kallingappa, Y Hou, M Annandale, K Mellor, ML Ward, CJ Barrett, DJ Crossman

## Abstract

Collagen VI has recently been strongly linked to poor outcomes in heart failure with preserved ejection fraction through increased endotrophin, a collagen VI-derived signalling molecule linked to fibrotic remodelling in cardiovascular disease. The mutation of collagen VI can result in Ullrich congenital muscular dystrophy and Bethlem myopathy, pointing to critical function in muscle physiology. However, the role of collagen VI in the heart is poorly understood. In human heart failure with reduced ejection fraction, collagen VI is increased within the remodelled T-tubules, suggesting a possible role in tubular structure and Ca^2+^ dynamics. To test this hypothesis, a global knockout of the collagen VI alpha 1 gene (Col6a1^-/-^) was generated in the rat. T-tubule structure and ryanodine receptor cluster organisation were unchanged, but echocardiography demonstrated reduced systolic function. Paradoxically, isolated cardiomyocytes from the Col6a1^-/-^ rat had increased Ca^2+^ transient amplitude and increased sarcoplasmic reticulum Ca^2+^ load. β-adrenergic stimulation further increased Ca^2+^ transient amplitude and was associated with diastolic Ca^2+^ release events in Col6a1^-/-^ cardiomyocytes. The disturbed Ca^2+^ dynamics are remarkedly similar to defects found in cardiac myocytes of the MDX mouse that has non-functional dystrophin protein. Together, these data suggest that collagen VI contributes to regulation of Ca^2+^ signalling in the heart through its linkage to the dystrophin-glycoprotein complex.

## Introduction

We previously identified increased levels of collagen VI within the remodelled T-tubules of the failing human heart with reduced ejection fraction (HFrEF), leading us to propose that collagen VI contributes to diminished Ca^2+^ signalling seen in this cardiac pathology^1^. Blunted and dyssynchronous Ca^2+^ handling dynamics in cardiac myocytes are strongly linked to the loss of T-tubules after experimental disruption^2–4^ and as a consequence of heart failure^5,6^. Collagen VI is rarely studied in the heart but is a focus of muscular dystrophy research, as mutations in this gene result in Ullrich congenital muscular dystrophy and Bethlem myopathy^7^. The function of collagen VI in striated muscle has been aided by the development of the global knockout of the collagen VI alpha 1 gene (Col6a1^-/-^) in the mouse as a model for Bethlem myopathy^8^. This animal demonstrated skeletal myopathy features on histology that included reduced fibre size and areas of necrosis but was otherwise phenotypically mild^8^. Paradoxically, the absence of functional collagen VI in the Col6a1^-/-^ mouse protected the heart from loss of function after myocardial infarction^9^. This was identified due to the absence of MI-induced fibrosis in the Col6a1^-/-^ heart. Subsequently, it was discovered that the proprotein of the Col6a3 gene contains a signalling peptide called endotrophin that drives fibrosis^10,11^ and contributes to the pathology of cancer, kidney disease, liver disease and cardiovascular disease^12^. For example, circulating levels of endotrophin are independently associated with poor outcomes in patients with HFrEF and heart failure with preserved ejection fraction (HFpEF)^13^. Most notably, plasma levels endotrophin provided a stronger predictor of outcomes in HFpEF patients than N-terminal pro B-type natriuretic peptide^13^. Yet very little is known about the function of collagen VI in the heart.

In order to increase our understanding of the function of collagen VI in the heart, we developed a CRISPR/Cas9 global knockout of the Col6a1 gene in the rat. Assessments included echocardiography to measure global heart function and dynamic investigation of Ca^2+^ transients and Ca^2+^ sparks in isolated cardiomyocytes. In addition, STimulated Emission Depletion (STED) microscopy of fixed ventricular tissue and isolated cardiomyocytes was used to measure the ultrastructure of the T-tubules and the cluster organisation of the cardiac calcium release channel, the ryanodine receptor (RyR2). These measures were used to test the hypothesis that collagen VI is involved in cardiomyocyte Ca^2+^ signalling in the heart.

## Methods

### Ethics Statement

All experimental procedures were approved by the Animal Ethics Committee of the University of Auckland (UoA) (No. AEC23882) in accordance with the New Zealand Government Animal Welfare Act.

### Animal Husbandry

Wistar IGS (Crl:WI) and Sprague Dawley [SD; Crl:CD(SD)] IGS rats were sourced from Charles River Laboratories, USA, and housed at the Vernon Jansen Unit, UoA. The rats were maintained in rooms with controlled temperature (22 ± 2°C), humidity (55 ± 10%), and a 12-hour light cycle (6:30 AM to 6:30 PM). Animals had ad libitum access to standard pellet food and 0.22µm filtered water.

### Generation of COL6A1 Knockout Rats

The generation of COL6A1 knockout (KO) rats was performed following a modified version of the procedure described^14,15^. Recombinant Cas9 protein (Alt-R™ S.p. Cas9 Nuclease 3NLS) was purchased from Integrated DNA Technologies (IDT), Singapore. Guide RNA (gRNA) was synthesized using the Precision gRNA Synthesis Kit (Thermo Fisher Scientific, USA) with DNA templates. CRISPR RNA (crRNA) design was carried out using the CCTOP software, and the sequence TAGATGCGGTCAAGTACTTC was selected. Full-length synthetic gRNA, “Alt-R CRISPR-Cas9 sgRNAs”, ordered from IDT consisted of a 100-base RNA oligonucleotide with the crRNA and trans-activating crRNA (tracrRNA) regions, including chemical modifications to enhance stability and nuclease resistance. The gRNA was diluted in microinjection buffer (1 mM Tris, 0.1 mM EDTA, pH 7.5) and stored at −20°C until use.

### Microinjection and Electroporation

The Cas9-gRNA ribonucleoprotein (RNP) complex was prepared by diluting gRNA and Cas9 to final concentrations of 50 ng/μl and 100 ng/μl, respectively. Wistar male rats (>10 weeks old) provided sperm, while Wistar females (4–5 weeks old) were used as oocyte source. Pseudopregnant SD females (8–16 weeks old) served as embryo recipients, and vasectomized SD males (>10 weeks old) were used to induce pseudopregnancy. Female Wistar rats were superovulated by intraperitoneal injection of pregnant mare serum gonadotropin (30 IU, ProSpec, Israel), followed 48 hours later by human chorionic gonadotropin (25 IU, Sigma-Aldrich, USA). Superovulated females were paired with male Wistar rats (1:1 ratio) overnight. The following day, pronuclear-stage embryos were collected from the oviducts and cultured in M2 medium (Sigma-Aldrich, USA) before being transferred to mR1ECM medium (Cosmo Bio Co. Ltd., Japan) for electroporation.

Zygotes with visible pronuclei were electroporated using a modified version of the procedure described by Koneko (2017)^16^. Electroporation was carried out with the NEPA21 electroporator to introduce the Cas9-gRNA ribonucleoprotein (RNP) complex into the one-cell stage embryos. Following electroporation, zygotes were cultured in mR1ECM medium (Cosmo Bio Co. Ltd., Japan) until transfer.

### Embryo Transfer and Genotyping

Healthy embryos were surgically transferred into the oviducts of pseudopregnant SD females mated with vasectomized males. Embryo transfer was performed either on the same day or the day following electroporation. Pups were delivered 20–21 days post-transfer. At weaning (3 weeks of age), ear punches were collected for DNA extraction. Genotyping was done via Sanger sequencing to confirm the presence of COL6A1 knockouts. Founder rats were backcrossed for three generations. Male Col6a1^-/-^ of 14-16 weeks of age were used for experimentation. Routine genotyping of the breeding colony was performed with qPCR (TransnetYX, USA).

### Echocardiographic functional Assessments

14-16-week-old male rats underwent echocardiographic assessments under general anaesthesia (2% isoflurane in oxygen) to ensure assessments using the VEVO-LAZR-X 3100 with a MX250 probe (14-28 Hz) linear array transducer coupled with digital ultrasound system (FUJIFILM Visual Sonics, Canada). Echocardiographic images were acquired as previously described^17^. All analyses were performed by an investigator blinded to study group allocation.

### Tissue Collection and Preparation

14-16-week-old male rats were anaesthetised using 2% isoflurane in 100% O_2_ as a carrier gas. Once unconscious, the rat was euthanised and underwent cervical dislocation. Immediately after euthanasia, the thoracic cavity was opened, and the heart was ligated and excised at the aorta, then placed in an ice-cold PBS buffer. The hearts were given slight compressions to release any remaining blood from within the ventricles and the coronary system. They were weighed after removing fat, excess connective tissue, and blood. Fixation of ventricular tissue blocks was carried out with 1% paraformaldehyde in phosphate-buffered saline (PBS) for 1 hour at 4°C, and then snap frozen with optimum cutting temperature (OCT) compound in 2-methyl butane using liquid nitrogen and stored in −80°C until cryosectioning for immunohistochemistry.

### T-tubule labelling of tissue

Left ventricular (LV) cross-sections 10 µm thick were cut using a Cryostat (CM3050, Leica, Germany) and mounted onto poly-L-lysine coated glass coverslips (22 × 50 mm, no. 1.5, Menzel™, Thermo Fisher Scientific, USA), which were taped on to labelled 25 × 75 × 1 mm microscope glass slides and stored at -80°C. For T-tubule labelling tissue sections were permeabilised in 1% Triton x-100 in PBS for 15 min and then blocked in Image-iT® FX signal enhancer (Thermo Fisher Scientific, USA) for 60 min at room temperature (RT). The sections were incubated with Wheat Germ Agglutinin Alexa Fluor 594 (Thermo Fisher Scientific, USA) diluted 1:100 in incubation buffer containing 1%BSA, 0.05% Triton x-100, 0.05% NaN3 in PBS. After incubation, the sections were washed and mounted in ProLong Gold Antifade Reagent (Thermo Fisher Scientific, USA).

### Cell isolation

The following cell isolation protocol was identical for both intracellular Ca^2+^ transient and diastolic Ca^2+^ spark experiments. Rats were anaesthetised using 2% isoflurane in 100% O_2_ as a carrier gas. Once unconscious, the rat was euthanised by cervical dislocation and hearts were excised at the aorta and placed in ice cold Ca^2+^ free Tyrode’s buffer. Ca^2+^ Free Tyrodes solution contained (in mM): 140 NaCl, 4 KCl, 10 Hepes, 1 MgCl2.6H2O, 10 Glucose, adjusted to pH 7.4 with 5 M NaOH. Immediately after excision, hearts were cannulated by the aorta, secured with a suture, and Langendorff-perfused with oxygenated Ca^2+^ free Tyrode’s buffer for 5 min at 37°C using a gravity fed system. During this time, the blood was cleared from the coronary circulation. Perfusion was then switched to 0.2 mM Ca^2+^ Tyrode’s containing enzymes: 275 U/mL Type 2 Collagenase (Worthington Biochemical Corp, USA) and 1.8 U/mL Protease (Sigma Aldrich, USA). After 9–12 min of enzymatic digestion, ventricles were cut off and immersed in 0.15 mM Ca^2+^ Tyrode’s containing 0.1% BSA (Sigma Aldrich, USA) and 0.05% Trypsin inhibitor (Worthington Biochemical, USA). The ventricles were minced to yield isolated, quiescent myocytes and extracellular Ca^2+^ was gradually increased to 1 mM.

### Loading of Ca^2+^ indicators

Once cells were isolated and increased to 1 mM Ca^2+^ Tyrode’s, they were divided into aliquots for the different measurements carried out. For intracellular Ca^2+^ transients, cells were loaded with 10 µM Fura-2/AM (Thermo Fisher Scientific, USA) dissolved in 20 µL dimethyl sulphoxide anhydrous (DMSO, ThermoFisher USA) with 20% pluronic (Thermo Fisher Scientific, USA) for 20 min at room temperature. Cells were then washed with 1 mM Ca^2+^ Tyrode’s solution for at least 10 min prior to imaging. For diastolic Ca^2+^ spark measurements, the same protocol was followed, but cells were loaded with 5 µM Fluo-4 (Thermo Fisher Scientific, USA) and then washed with 1.8mM Ca^2+^ Tyrode’s solution.

### Intracellular Ca^2+^ transient measurements

Upon loading with Fura-2, cardiomyocytes were transferred to a cell bath, field stimulated at 1 Hz (room temperature) and continuously super-perfused with 1 mM Ca^2+^ Tyrode buffer. Myocytes were imaged using a 20x fluorescent objective lens (0.75 NA) and illuminated with alternating 340 nm and 380 nm excitation wavelengths every 5 ms using an Optoscan monochromator and spectrofluorometric, PMT-based system (Cairn, UK). Emitted 510 nm ± 15 nm fluorescence was acquired at 400 Hz using Acquisition Engine Software (Cairn, UK) from the whole cell. The total emitted fluorescence provided a measure of cytosolic [Ca^2+^] fluxes. The cardiomyocytes were subjected to interventions used to modulate different mechanisms of EC coupling. The first intervention investigated was the response to different stimulation frequencies. Myocytes were subjected to periods of 0.2 Hz, 0.5 Hz and 1 Hz stimulation until steady state was achieved at each frequency. Once myocytes were contracting at steady state (1 Hz), both the flow and the stimulus were switched off and a bolus of 20 mM caffeine was applied to the bath. Once a large caffeine induced Ca^2+^ release was observed, caffeine was washed off and a 1 Hz stimulus was re-commenced. As the myocytes returned to steady state, super-perfusion with 1 mM Ca^2+^ Tyrode’s containing 1 µM isoproterenol (ISO) was initiated to determine the response to β-adrenergic stimulation. Intracellular Ca^2+^ transient data from loaded cardiomyocytes were subsequently analysed using a custom-written IDL program (IDL version 6.2, Research Systems Inc., USA) to determine the various Ca^2+^ transient parameters.

### Ca^2+^ spark measurements

Cardiomyocytes loaded with Fluo-4 in 1.8 mM Ca^2+^ Tyrode’s were transferred to a cell bath fixed above a 40x water-immersion objective lens (NA 1.1) to the stage of an inverted laser scanning confocal microscope (LSM 800, Zeiss, Germany). Prior to recording, cells were continuously stimulated for at least 20 s, and the objective was focused on the surface of a single, healthy, rod-shaped cardiomyocyte. The stimulus was then stopped, and a snapshot of the myocyte was taken, which was used as a guide for placement of the line-scan along the cell. Line scans were always placed along the longitudinal axis of the cell, avoiding nuclei. For Ca^2+^ spark measurements, line-scan images were recorded bi-directionally in quiescent myocytes at approximately 2.4 ms/line with a 0.1-0.15 µm pixel resolution. Images were obtained across a 4 s rest period after 20s of continuous 1 Hz stimulation, while line-scan images were also obtained during 1 Hz stimulation of the cardiomyocyte, to confirm that the cell was healthy and responsive to stimuli (i.e., the presence of spatially uniform Ca^2+^ transients).

Ca^2+^ spark data from the line-scan images were subsequently analysed using the SparkMaster algorithm plugin on Image J FIJI^18^ to reduce investigator error and bias. SparkMaster parameters were set to the scanning speed (i.e., 2.4 ms/line) and pixel size (0.1 µm) of the line scan. Sparks were analysed using “5” for the number of intervals and criteria of “3.8”. This criterion indicated that the detection of events was 3.8 times the standard deviation of background noise, ensuring high sensitivity and reduced likelihood of false-positive detections^18^. The output from the SparkMaster detection determined multiple Ca^2+^ spark parameters (see Fig. 5).

### Cell fixation

Upon isolation of cardiomyocytes, a cell suspension in low Ca^2+^ Tyrode’s were centrifuged for 2 minutes at 700 rpm. The supernatant was removed and the pellet was resuspended in 2 % PFA in PBS and incubated at room temperature for 10 minutes. The tubes were then centrifuged for 2 minutes at 1100 rpm, and the supernatant was removed and the pellet resuspended in PBS. After at 10 min incubation, cells were centrifuged again, and the supernatant was replaced with a storage solution containing 0.5 % BSA and 0.05 % Na+ azide in PBS. The cells were stored at 4°C until labelling.

### RyR2 labelling of isolated myocytes

Previously fixed cells were resuspended, an aliquot was centrifuged for 3 minutes at 3000 rpm, and the supernatant discarded, a procedure repeated between each of the following steps. Cells were then incubated in a permeabilisation solution containing 1% triton in PBS for 15 minutes at RT. Cells were then incubated in Image-iT FX signal enhancer for 30 minutes at RT. Cells were then incubated with the primary antibody Mouse anti-RyR (MA3-916, Thermo Fisher Scientific, USA) that was diluted 1:100 in incubation solution (1%BSA, 0.05% Triton x-100, 0.05% NaN3 in PBS) overnight at 4°C. The following day, cells were washed 3 times in PBS and resuspended in a solution containing the secondary antibody Goat anti-mouse Abberiror STAR RED (Abberior, Germany) diluted 1:200 and incubated for 2 hours at RT. Cells were washed in PBS and mounted in ProLong Gold mounting medium(Thermo Fisher Scientific, USA) on a glass slide and poly-L-lysine coated #1.5 coverslips.

### STED Microscopy

Two-dimensional STED images of LV sections labelled for T-tubules/WGA (Alexa Fluor 597) or ventricular myocytes labelled for RyR2 (Abberior Star Red) were obtained using an Olympus IX83 Abberior Facility Line STED microscope (Abberior Instruments, Germany) at the Biomedical Imaging Research Unit, University of Auckland, equipped with a 63x 1.4 NA oil immersion objective. Before each imaging session, the STED microscope was allowed to warm up, and routine alignment checks were conducted using Autoalignment Slide (Abberior Instruments, Germany). All images were collected from longitudinally orientated cardiomyocytes. For imaging of T-tubules (Alexa Fluor 597) the pixel size was set to 20 × 20 nm, the excitation laser 561 nm, pinhole 1 Airy unit, STED laser 775 nm, detector 588-698 nm, line accumulation of 3, and STED gating of 750 ps (+8 ns). For the imaging of RyR2 (Abberior Star Red) the pixel size was set to 10 × 10 nm, the excitation laser 640 nm, pinhole 0.8 Airy unit, STED laser 775 nm, detector 650-755 nm, line accumulation of 3, and STED gating of 750 ps (+8 ns).

### Image analysis

T-tubule images were deconvolved with Huygens Essential version 24.04 (Scientific Volume Imaging, The Netherlands, http://svi.nl). T-tubule images were then analyzed for the following metrics: regularity (T-power) and T-tubule skeleton area as previously described^19^. This involved using the pixel classification feature of the machine learning software, Ilastik version 1.4 to segment WGA labelling into both a cellular binary mask of the myocyte area and a binary mask of the T-tubules. The sarcomeric regularity of the T-tubule masks was assessed in the frequency domain using custom ImageJ/FIJI macros. The T-tubule mask was then converted into a 1-pixel-wide skeleton. T-tubule length was estimated for each cell by normalizing the length (area) of the skeleton by cell area and expressing it as a percentage. T-tubule diameter was calculated as previously described^20^. This involved using ImageJ/FIJI and extracting a line profile perpendicular to the T-tubule axis and then fitting a Gaussian function to the data and calculating the Full Width Half Max (FWHM) as a measure of diameter. RyR2 images were first processed with the PureDenoise algorithm^21^ followed by the rolling ball algorithm to remove the background^22^ and then converted to binary mask with the isodata algorithm^23^ using custom macros written in ImageJ/FIJI. RyR2 cluster sizes, cluster centroid-to-centroid distances, and cluster edge-to-edge distances were measured using Python code utilising the scipy.ndimage library as previously described^24,25^.

### Statistics

Values presented are means ± SEM. T-tests were used to test for differences in morphometric and echocardiography. For image and Ca^2+^ measurements, a linear mixed model analysis was used with a two-level hierarchy (level 1: group (wildtype vs. Col6a1-/-) and level 2: random factor animal (myocytes per animal). Statistical analyses were carried out using IBM SPSS Statistics (Version: 28.0.1.1(15)) with P<0.05 considered significant. Data are available from authors upon request.

## Results

The appearance of the freshly excised Col6a1^-/-^ rat heart was unremarkable and similar to hearts from the wildtype animals (Fig.1a). However, heart wet weights were significantly lower in the Col6a1^-/-^ animals (Fig. 1b). Likewise, both body weight and tibia length were significantly reduced in comparison to equivalent-age wildtype littermates (Fig. 1c-d). When the heart weight was normalised to body weight, the heart was significantly enlarged, but no difference between groups was observed when the heart weight was normalised to tibia length (Fig. 1e-f). Previously our group has shown collagen VI is present in the T-tubules of the heart^1^. We therefore sought to determine if the absence of collagen VI induced structural and functional effects on the heart of Col6a1^-/-^ rats. T-tubule structure was assessed by using STED super-resolution microscopy to image tissue sections labelled with wheat germ agglutinin (WGA, Fig. 1i). The STED imaging showed no difference between wildtype and Col6a1^-/-^ T-tubule structure as assessed by the metrics, T-power (regularity), T-tubule length, and T-tubule diameter (Fig. 1i-k).

**Fig. 1.**
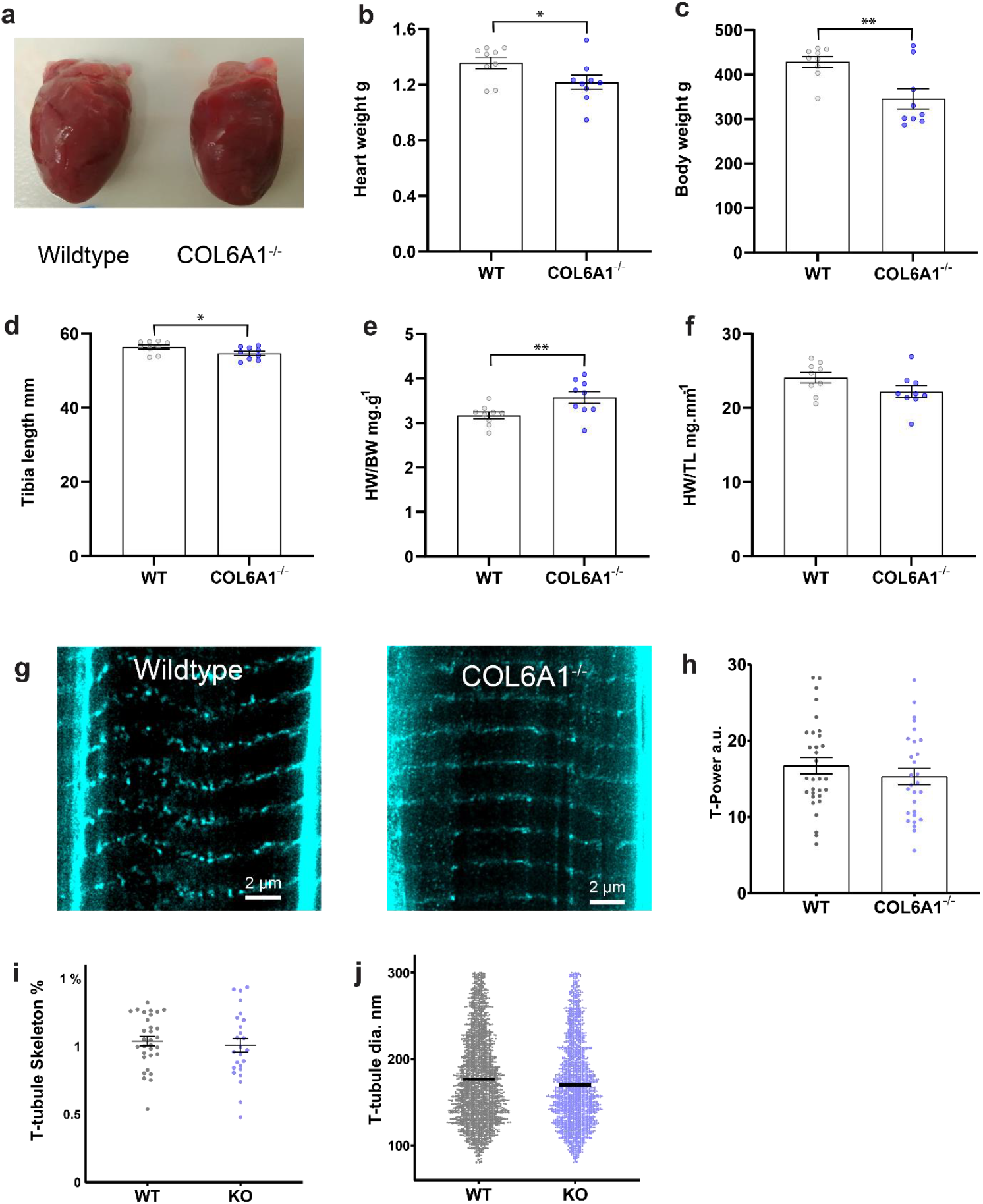
The absence of functional collagen VI in the Col6a1^-/-^ rat results in reduced body weight relative to wildtype animals but normal heart structure at the anatomical and cellular levels. **a** Example images of the wildtype and Col6a1^-/-^ rat heart. **b** Heart weight. **c** Body weight. **d** Tibia length. **e** Heart weight normalised to body weight (HW/BW). **f** Heart weight normalised to tibia length (HW/TL). **g** Example images of STED microscopy of WGA, labelled T-tubules in wildtype and Col6a1^-/-^ rat heart. **h** T-tubule frequency analysis (T-power). **i** T-tubule length (Skeleton %). **j** T-tubule diameter. For panels, **b-f** data are mean ± SEM with each point representing a single animal, n = 9 and n = 9 biologically independent replicates for wildtype and Col6a1^-/-^ rats, respectively. Statistical significance was determined with a t-test. For panels, **i-j** data are mean ± SEM with each point representing a single myocyte, n = 5 and n = 5 biologically independent replicates for wildtype and Col6a1^-/-^ rats, respectively, with 4-9 cellular replicates for each heart. Mixed model analysis with two-level hierarchy (level 1: group (wildtype vs. Col6a1^-/-^) level 2: random factor animal (4-9 myocytes per animal). *=p<0.05, **=p<0.01, ***=p<0.001.

Echocardiography assessment of Col6a1^-/-^ rat showed a decrease in systolic function (Fig. 2). M-mode echocardiography showed a significantly reduced ejection fraction, fractional shortening, stroke volume, and cardiac output in the Col6a1^-/-^ rat (Fig. 2a-e). This was not related to heart rate as there was no difference in heart rate during echocardiology measurements (Fig. 2f). However, normalisation of stroke volume and cardiac output to body weight showed no significant change in stroke volume or cardiac output (Fig. 2G-H). Pulse wave Doppler imaging showed a significant increase in the ratio of early (E) to late (A) mitral valve blood flow velocity (E/A ratio) in Col6a1^-/-^ rat compared to wildtype littermates (Fig. 2i-j). This finding often indicates restrictive filling but can also result from reduced atrial function^26^. The increased E-wave deceleration time indicates that the ventricle does not have increased stiffness as would be expected with restrictive filling (Fig. 2k). However, the decreased A-wave velocity (atria contraction) compared to the unchanged E-wave velocity (ventricular relaxation) indicates the elevated E/A ratio in the Col6a1^-/-^ rat is due to impaired atrial contraction (Fig. 2l-m).

**Fig. 2.**
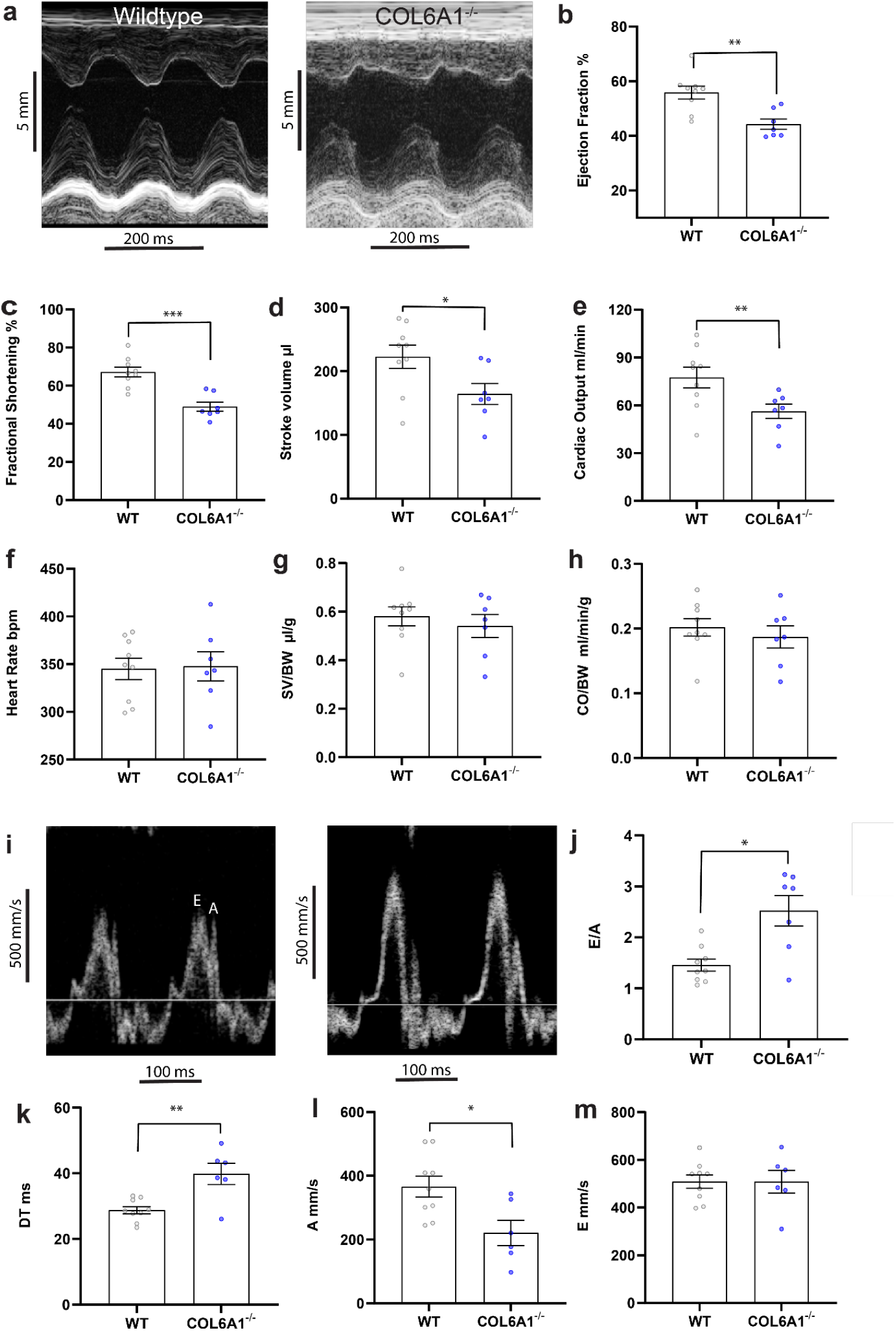
Echocardiography assessment of Col6a1^-/-^ rat showed impaired systolic function. **a** Example M-mode traces of the wildtype and Col6a1^-/-^ rat heart. **b** Ejection fraction. **c** Fractional shortening. **d** stroke volume. **e** cardiac output. **f** Heart rate. **g** Stroke volume normalised to body weight (SV/BV). **h** Cardiac output normalised to body weight (CO/BW). **i** Example pulse-wave Doppler traces of the wildtype and Col6a1-/-rat heart. **j** E wave over A wave ratio (E/A). **k** E wave deceleration time. **l** A wave velocity. **m** E wave velocity. data are mean ± SEM with each point representing a single animal, n = 9 and n = 6-7 biologically independent replicates for wildtype and Col6a1^-/-^ rats, respectively. Statistical significance was determined with a t-test. *=p<0.05, **=p<0.01, ***=p<0.001.

To determine if the reduced contractile function of the heart was due to changes in excitation-contraction coupling, the response of isolated ventricular myocytes to electrical stimulation was recorded following fura-2/AM loading (Fig. 3). Representative Ca^2+^ transients (340/380 fura-2 ratio) for the Col6a1^-/-^ and wildtype rats are shown in Fig 3a. Analysis showed the Col6a1^-/-^ rat had increased Ca^2+^ transient amplitude (peak systolic – diastolic) relative to wildtype rat at 1 Hz stimulation (Fig. 3b). Increasing stimulation frequency (0.2, 0.5, 1 Hz) increased the Ca^2+^ transient amplitude in both the Col6a1^-/-^ and wildtype rat cardiomyocytes, with a greater extent of increase evident in the Col6a1^-/-^ cells (Fig 3b). The Ca^2+^ transient maximum rate of rise was significantly increased at 1 Hz vs 0.5 and 0.2 Hz in Col6a1^-/-^ cardiomyocytes, an effect that was not observed in wildtype cells (Fig. 3c). There was no difference in the time constant of decay (tau) between the groups but stimulation frequency significantly reduced tau in both Col6a1^-/-^ and wildtype rat cardiomyocytes (Fig. 3d). There was no change in time to peak Ca^2+^ transient between groups or with stimulation frequency (Fig. 3e). In addition, there was no difference in diastolic Ca^2+^ between groups, but stimulation frequency significantly increased diastolic Ca^2+^ in both Col6a1^-/-^ and wildtype rat cardiomyocytes (Fig. 3f).

**Fig. 3.**
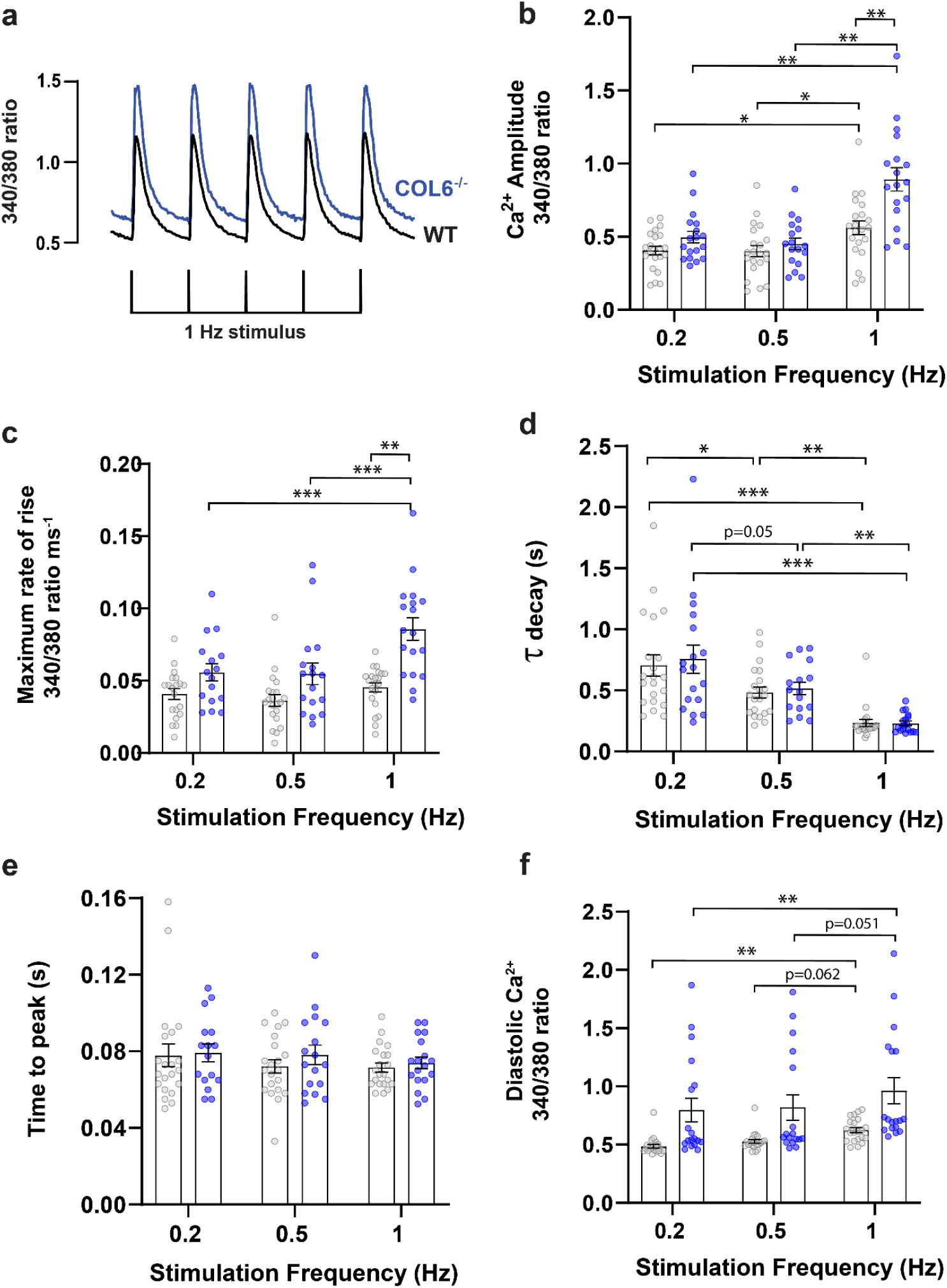
Col6a1^-/-^ rat cardiac myocytes have increased fura-2 Ca^2+^ transient that is associated with increased sarcoplasmic reticulum Ca^2+^ load. **a** Exemplar fura-2 Ca^2+^ transient showing an increase in the Col6a1^-/-^ rat at 1 Hz stimulation. **b** Peak Ca^2+^ transient amplitude at 0.2, 0.5, and 1 Hz stimulation frequency. **c** Maximum rate of rise of Ca^2+^ transient. **d** Ca^2+^ transient decay, Tau (τ). **e** Time to peak Ca^2+^ transient amplitude. **e** Diastolic Ca^2+^ concentration. Data are mean ± SEM with each point representing a single myocyte, n = 4 and n = 5 biologically independent replicates for wildtype and Col6a1^-/-^ rats, respectively, with 3-8 cellular replicates for each heart. Mixed model analysis with two-level hierarchy (level 1: group (wildtype vs. Col6a1^-/-^) level 2: random factor animal (3-9 myocytes per animal). *=p<0.05, **=p<0.01, ***=p<0.001.

To determine the response to β-adrenergic stimulation, cardiomyocytes were electrically stimulated at 1 Hz and exposed to 1 µM isoprenaline. Isoprenaline significantly increased the Ca^2+^ transient amplitude in both wildtype and Col6a1^-/-^ cardiomyocytes, with a significantly larger response in the Col6a1^-/-^ cardiomyocytes (Fig. 4a). The Ca^2+^ transient rate of rise was increased after the addition of isoprenaline in the Col6a1^-/-^ cardiomyocytes and was significantly larger than the rate of rise in wildtype cardiomyocytes (Fig. 4b). Isoprenaline similarly decreased the time constant of the Ca^2+^ transient decay for both groups (Fig. 4c). Examination of the Ca^2+^ during isoprenaline stimulation revealed spontaneous diastolic Ca^2+^ release events in ∼50% of the Col6a1^-/-^ cardiomyocytes, which were not observed in wildtype myocytes (Fig. 4d). The caffeine-induced fura-2 Ca^2+^ release was then assessed to determine if increased sarcoplasmic reticulum (SR) Ca^2+^ levels could explain the elevated Ca^2+^ transients observed in the Col6a1^-/-^ cardiomyocytes. Cardiomyocytes were first stimulated at 1 Hz and once steady state was achieved stimulus was switched off and a bolus of 20 mM caffeine was applied to the bath. This analysis demonstrated a significant increase in the peak of the caffeine-induced Ca^2+^ transient in Col6a1^-/-^ cardiomyocytes, indicating increased SR Ca^2+^ load (Fig. 4e-f). There was an apparent decrease in Ca^2+^ decay that is indicative of the flux of Ca^2+^ via the Na^+^/Ca^2+^ exchanger, but this change did not reach significance (Fig. 4g).

**Fig. 4.**
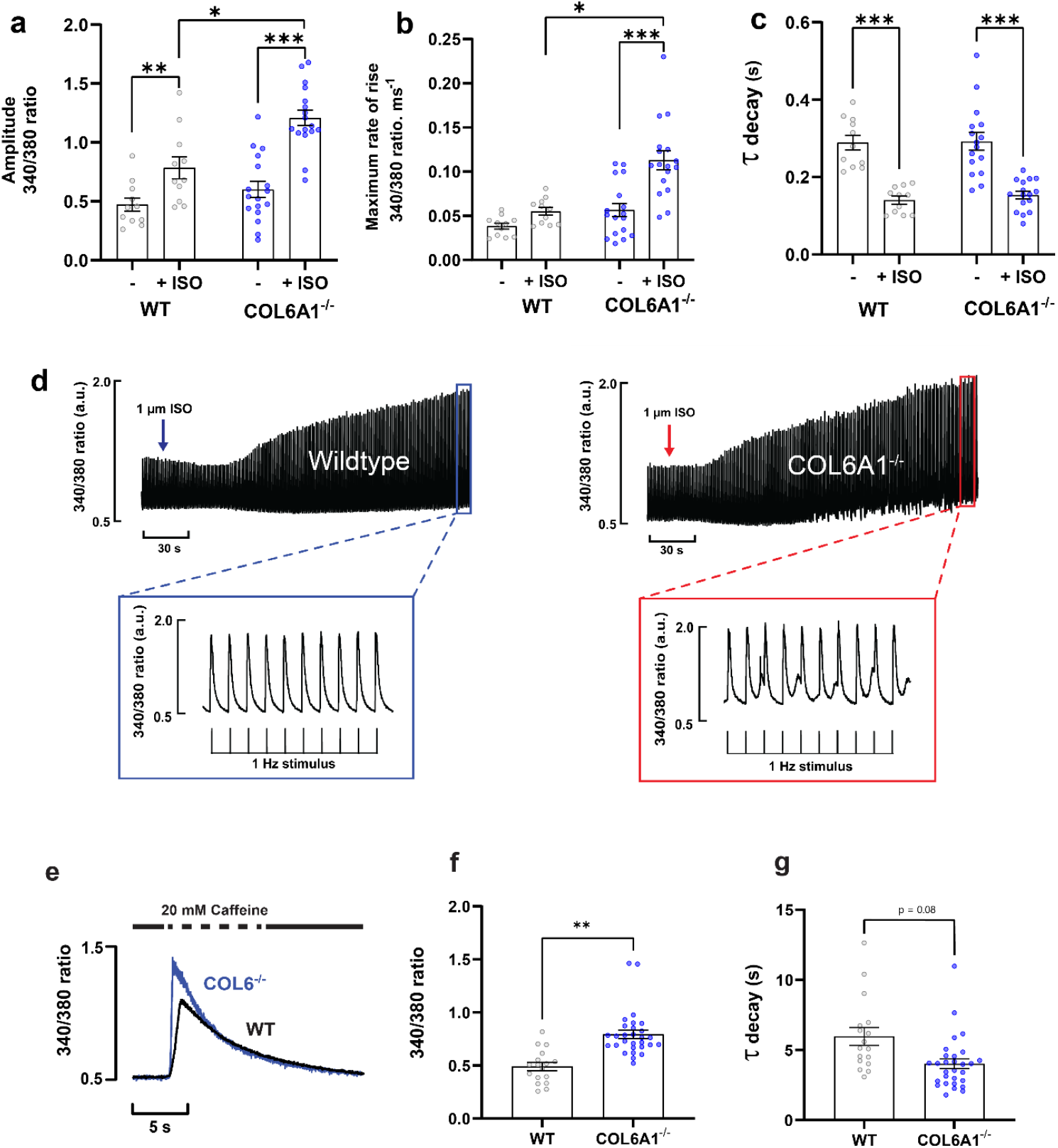
β-adrenergic stimulation elicits an increased Fura-2 Ca^2+^ transient in Col6a1^-/-^ rat cardiac myocytes. **a** Ca^2+^ transient amplitude in wildtype and Col6a1^-/-^ rat cardiac myocytes after the addition of isoprenaline (iso). **b** Ca^2+^ transient maximum rate of rise in wildtype and Col6a1^-/-^ rat cardiac myocytes after the addition of isoprenaline (iso). **c** Ca^2+^ transient decay (τ) in wildtype and Col6a1^-/-^ rat cardiac myocytes after the addition of iso. **d** example traces from wildtype and Col6a1^-/-^ rat cardiac myocytes stimulated at 1 Hz demonstrating the response to iso. Note the small diastolic peaks between the systolic peaks in the Col6a1^-/-^ rat cardiac myocyte. Data are mean ± SEM with each point representing a single myocyte, n = 3 and n = 5 biologically independent replicates for wildtype and Col6a1^-/-^ rats, respectively, with 1-6 cellular replicates for each heart. Mixed model analysis with two-level hierarchy (level 1: group (wildtype vs. Col6a1^-/-^) level 2: random factor animal (1-8 myocytes per animal). **e** Exemplar caffeine-induced fura-2 Ca^2+^ transient showing an increase in the Col6a1^-/-^ rat at 1 Hz stimulation. **f** Peak caffeine-induced Ca^2+^ transient amplitude. **i** Ca^2+^ transient decay, τ, after caffeine-induced release. Data as for Fig. 3. *=p<0.05, **=p<0.01, ***=p<0.001.

Confocal line-scanning of quiescent ventricular cardiomyocytes loaded with Fluo-4 was used to determine if changes in Ca^2+^ spark morphology could help explain the increased Ca^2+^ transient amplitude in the Col6a1^-/-^ rat cardiomyocytes. A representative line scan from a wildtype and Col6a1^-/-^ rat cardiomyocyte is shown in Fig. 5A. The data showed an apparent increase in spark frequency in the Col6a1^-/-^ myocytes, but this did not reach significance (Fig. 5b). The amplitude of the Ca^2+^ sparks was significantly reduced in the Col6a1^-/-^ myocytes (Fig. 5c). The apparent Ca^2+^ spark mass was reduced in the Col6a1^-/-^ myocytes, but this did not reach significance (p=0.09, Fig. 5d). The diastolic SR Ca^2+^ “leak” attributable to sparks (frequency x mass) was not different between the animal groups (Fig. 5e). RyR2 cluster dimensions were then examined. RyR2 cluster arrangement has been shown to correlate with spark dynamics in normal and failing cardiomyocytes ^27–29^. Representative STED images of RyR2 labelling in wildtype and Col6a1^-/-^ cardiomyocytes, and the binary masks used to measure the cluster parameters are presented in Fig. 5f-g. Analysis of RyR2 cluster parameters showed no change in RyR2 cluster size, RyR2 cluster nearest neighbour centroid to centroid distances, and RyR2 cluster nearest neighbour edge to edge distances (Fig. 5h-j).

**Fig. 5.**
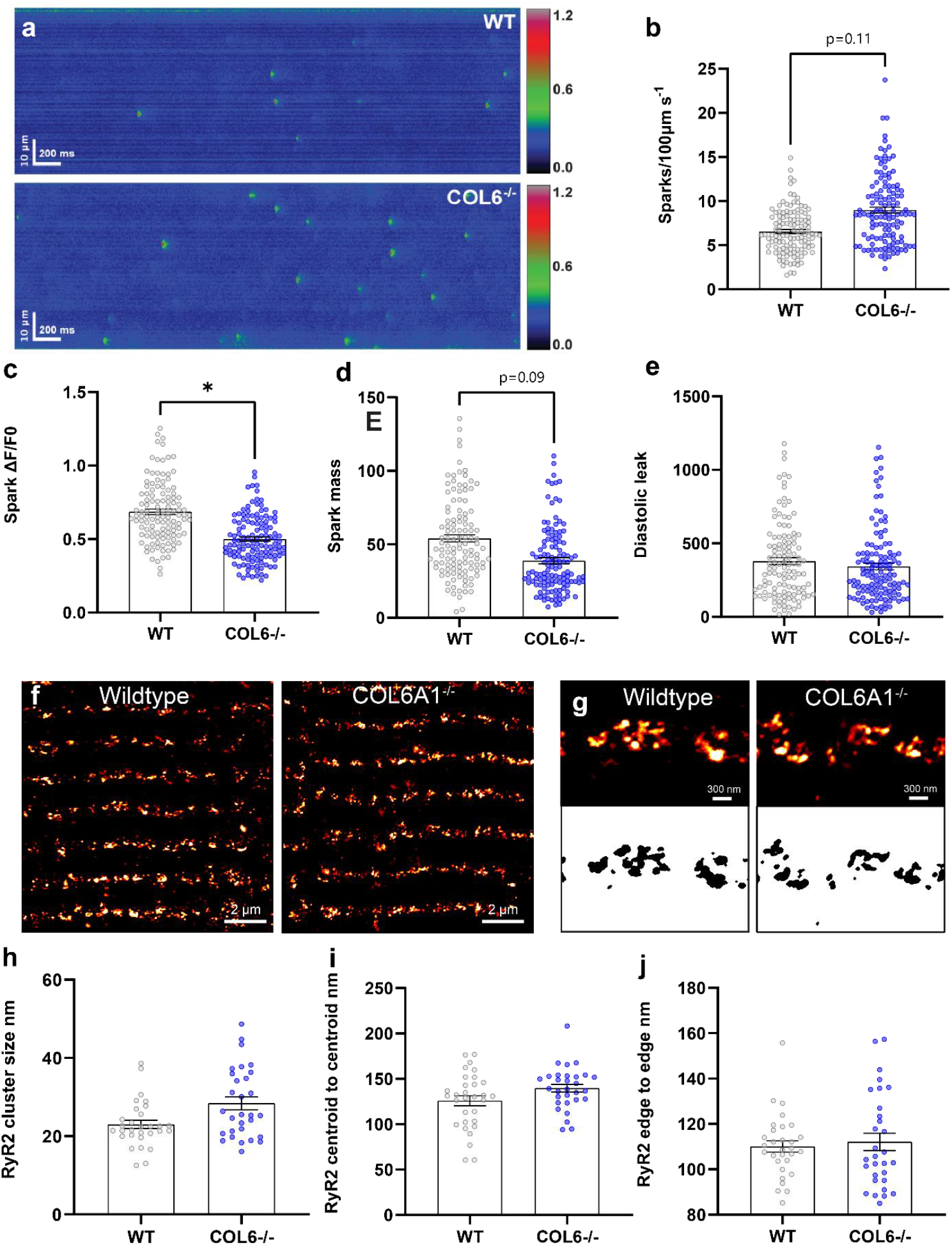
Quiescent Ca^2+^ sparks, and RyR2 cluster organisation do not explain the increased Ca^2+^ transient in Col6a1^-/-^ rat cardiac myocytes. **a** Examples of confocal line-scans from Fluo-4 loaded cardiac myocytes from wildtype and Col6a1^-/-^ rat. **b** Ca^2+^ spark frequency. **c** Ca^2+^ spark amplitude. **d** Ca^2+^ spark mass. **e** Ca^2+^ spark diastolic leak. **f** example STED microscopy images of cardiac myocytes from cardiac myocytes from wildtype and Col6a1^-/-^ rat. **g** enlarged RyR2 images (top panel) and corresponding binary mask used to measure RyR2 cluster parameters. **h** RyR2 cluster size. **i** RyR2 cluster nearest neighbour centroid to centroid distance. **j** RyR2 cluster nearest neighbour edge to edge distance. Ca^2+^ spark data are mean ± SEM with each point representing a single myocyte, n = 5 and n = 5 biologically independent replicates for wildtype and Col6a1^-/-^ rats, respectively, with 9-28 cellular replicates for each heart. Mixed model analysis with two-level hierarchy (level 1: group (wildtype vs. Col6a1^-/-^) level 2: random factor animal (9-28 myocytes per animal). RyR2 cluster data are mean ± SEM with each point representing a single myocyte, n = 5 and n = 5 biologically independent replicates for wildtype and Col6a1^-/-^ rats, respectively, with 5-8 cellular replicates for each heart. Mixed model analysis with two-level hierarchy (level 1: group (wildtype vs. Col6a1^-/-^) level 2: random factor animal (5-8 myocytes per animal). *=p<0.05, **=p<0.01, ***=p<0.001.

## Discussion

Echocardiographic data demonstrated that the absence of functional collagen VI in Col6a1^-/-^ rat is associated with reduced systolic function of the heart at ∼16 weeks old. This was demonstrated by reduced ejection fraction, fractional shortening, stroke volume, and cardiac output. Moreover, doppler measurements showed reduced atrial contraction. These data are consistent with our previous study, where we proposed that collagen VI is part of the dystrophin-glycoprotein complex (DGC)^1^. The DGC links the myocyte cytoskeleton to the extracellular matrix and is part of the myocyte costamere complex that is crucial for force production^30^ and can explain the reduced systolic function described here, particularly as muscular dystrophy patients often display cardiomyopathy^31^. Supporting this view, echocardiography demonstrates a loss of systolic function in the MDX mouse heart^31,32^ although this decline occurs after ∼25 weeks in the life course of the mouse. In contrast to our results, the cardiac ejection fraction is preserved in the Col6a1^-/-^ mouse heart at ∼26 weeks of age^9^ and cardiac function is described as normal or only mildly disturbed in collagen VI muscular dystrophy patients^33,34^. Notably, in the Col6a1^-/-^ rat, when stroke volume and cardiac output was normalised to body weight, there was no difference, suggesting changes are functionally mild.

Intriguingly, the absence of functional collagen VI increases the Ca^2+^ transient in Col6a1^-/-^ rat cardiomyocytes. The changes in Ca^2+^ handling were not due to structural changes as the T-tubules were found to be intact in the Col6a1^-/-^ rat heart, which would be expected to maintain or preserve Ca^2+^ handling. So what is causing the increased Ca^2+^ release? Changes in RyR2 cluster organisation can perturb Ca^2+^ release^27–29^. However, RyR2 cluster structure is unchanged in the Col6a1^-/-^ rat heart and the Ca^2+^ spark mass is also unchanged. The most likely explanation is the higher SR Ca^2+^ load that we documented in Col6a1^-/-^ cardiomyocytes, as greater SR Ca^2+^ load is known to increase Ca^2+^ release^35^. This increase in SR Ca^2+^ load also provides an explanation for the heightened response to β-adrenergic stimulation. Interestingly, under these conditions, spontaneous diastolic Ca^2+^ release events were seen in ∼50% of Col6a1^-/-^ rat myocytes, a feature that would be electrogenic and potentially trigger arrhythmias.

The changes in Ca^2+^ release observed in the Col6a1^-/-^ rat cardiomyocytes are remarkedly similar to Ca^2+^ release in cardiomyocytes from the MDX mouse^36^ that has a mutated and dysfunctional dystrophin^37,38^. The cardiomyocytes from the MDX mouse have elevated Ca^2+^ transients and SR Ca^2+^ stores^36^, in addition to *in vivo* ventricular arrhythmias when exposed to β-adrenergic stimulation^39^. In the present study, we demonstrated that the absence of collagen VI caused a loss of systolic function in the heart while at the same time increasing Ca^2+^ transient amplitude. These data support the hypothesis that collagen VI is involved in both force transduction and in regulating Ca^2+^ signalling within the heart.

## Acknowledgments

The authors acknowledge the microscope resources provided by the Biomedical Imaging Research Unit at the Faculty of Medical and Health Sciences at the University of Auckland. We thank Steph Lindsay for her work in caring for the animals, and acknowledge the AH Somerville Foundation whose generous donation funded the purchase of the ultrasound machine.

